# The expression of IL-35 in the prophase of liver failure and its preliminary exploration for the mechanism of immunoregulation by IL-35 and glucocorticoids

**DOI:** 10.1101/2025.01.23.634472

**Authors:** Yan Wang, Xin-chi Zhang, Yu-fan Xiong, Xin-yi Li, Xiao-ping Huang, Jing Gu, Wei Sun, Li Chen, Jian-he Gan

**Affiliations:** Department of Infectious Diseases, The First Affiliated Hospital of Soochow University, Suzhou, 215000, Jiangsu, China

**Author notes:** Corresponding author; Jian-he Gan. These authors have contributed equally to this work and share corresponding author (Li Chen, Jian-he Gan). ETHICS APPROVAL Approved by the Ethics Committee of The First Affiliated Hospital of Soochow University, compliant with principles of the Helsinki Declaration. E-mail : Yan Wang, Xin-chi Zhang, Yu-fan Xiong, Xin-yi Li, Xiao-ping Huang, Jing Gu, Wei Sun, Li Chen. FUNDING Funded, by National Major Science and Technology Projects for the 13th Five Year Plan (No.2017ZX10203201002-002), Tianqing Liver Disease Research Fund (TQGB20210134) and Suzhou Science and Education Xingwei Youth Science and Technology Project (KJXW2020003).

**Keywords:** IL-35, Glucocorticoids, Liver Failure

## Abstract

**Background and Aims:** To determine appropriate dosages for mice models in liver failure prophase and explore IL-35 and cytokine levels. Also, investigate changes in Treg/ Th17 ratio, IL-35, and IL-17 in patients with HBV-induced liver failure before and after treatment.

**Methods:** We induced liver failure in mice with different doses of concanavalin and assessed severity through serum biochemistry and pathology. Mice were divided into control, model, IL-35 plasmid, anti-IL-35, and dexamethasone groups. IL-35 and IL-17 expressions in liver tissue were evaluated via immunohistochemistry, fluorescence staining, and ELISA. Treg/Th17 ratio and IL-35 and IL-17 levels in peripheral blood of patients were measured using flow cytometry and ELISA, compared to 22 healthy individuals.

**Results:** The optimal dose for the liver failure model was 20 μ g/ kg concanavalin. The IL-35 plasmid and dexamethasone groups showed significantly lower serum TBil, ALT, AST, IL-4, IL-17, TNF-α , and liver histopathology scores compared to the model group, while the anti-IL-35 group showed higher levels (P<0. 05). In HBV-PLF patients, TBil, ALT, and AST significantly decreased, and PTA increased after glucocorticoid treatment (P<0. 05). Treg/ Th17 ratio was lower in HBV-PLF patients compared to the healthy group (P<0. 05), with higher IL-35 and IL-17 levels (P<0. 05). Post-treatment, Treg increased and Th17 decreased significantly; IL-17 and Treg/ Th17 ratios increased, while IL-35 decreased (P<0.05). IL-35 positively correlated with the Treg/Th17 ratio.

Conclusions

20 μ g/ kg concanavalin proper dose for mice model. IL-35 may protect in liver failure. Glucocorticoids help maintain immune balance, prevent failure, increase ratios and IL-35 in HBV-PLF patients, and IL-35 related to Treg/Th17 balance.

## Introduction

Liver failure is a clinical syndrome characterized by extensive degeneration and necrosis of hepatocytes within a short time due to viral infections, drug exposure, or other toxins [1,2]. The mortality rate associated with liver failure is high, exceeding 60%, While liver transplantation remains the only viable option for many patients, the availability of suitable donors is limited, and the procedure can be costly, making it a realistic option for a select few patients only [3]. Therefore, it is crucial to identify effective interventions to prevent the progression of liver failure during the prophase of liver failure (PLF)[4].

Regulatory T cells perform a vital function in maintaining immune homeostasis within the body by reducing both the innate and adaptive immune responses. IL-35 plays a crucial role in mediating the immunosuppressive activities of Treg cells [5–6]. Our previous findings revealed that overexpression of IL-35 in patients with liver failure increased the risk of developing secondary infections, while a deficiency in the expression of IL-35 was associated with poor prognosis [7–9]. Therefore, in our study, we want to know more about the role of IL-35 in the progresion of prophase of liver failure.

## Methods

### Mouse Models

We used female BALB/c mice weighing 20-25g and aged between eight to twelve weeks, purchased from Zhaoyan (Suzhou) New Drug Research Center Co., Ltd, for our study. After inducing the liver failure model, mice from each experimental group were humanely euthanized using cervical dislocation, and blood was collected by orbital blood collection. The liver of each mouse was quickly removed, and the left lobe was set aside for use while the rest of the liver tissues were cut into approximately 2mm thick pieces with a size of about 1cm. These pieces of liver tissue were then soaked and fixed in prepared formalin solution for 48 hours. The remaining liver tissue was stored at a temperature of -80°C in the refrigerator for further research. All animal procedures were conducted following the guidelines established by the National Institutes of Health for the care and use of laboratory animals and were approved by the institutional review committee of Soochow University.

### Biochemical assessments

In the mice, we measured serum total bilirubin, alanine aminotransferase (ALT), and aspartate aminotransferase (AST) levels. To do this, we allowed the samples to stand at room temperature for 1 hour and then centrifuged them at 3000 R/ min for 20 minutes at 4 °C. The resulting serum was sent to our hospital for biochemical detection of TBIL, ALT, and AST using a Hitachi 7500 automatic analyzer (Hitachi, Tokyo, Japan). The results were expressed in U/ L.

### HE staining of mouse liver tissue

The liver samples were fixed in 10% neutral buffered formalin, processed, and embedded in paraffin. The embedded samples were then sliced into 5μm sections and stained with hematoxylin and eosin (H& E) to visualize the histological changes in the liver tissue.

### Immunohistochemistry and Immunofluorescence for IL-35, IL-17, Treg

To evaluate changes in inflammatory mediators in the liver tissue, we stained paraffin-embedded liver tissue sections using antibodies against Anti-IL-12A (Abcam, USA), anti-IL-17A (Abcam, USA), and anti-FOXP3 (Affinity, USA). The staining process was performed using a polymer detection system for double immunohistology staining (Zhong Shan Gold Bridge Biotechnology, Beijing, China). We employed a semiquantitative scoring system to assess the histological changes after the application of HRP-labeled broad-spectrum secondary antibody (or fluorescent secondary antibody) on the sample slides, which were then incubated in a wet box.

### Flow cytometry for Th17 and Treg

To assess the phenotype of immune cells in the liver tissue, we harvested cells and transferred them to FACS tubes. The cells were then stained with anti-CD4-PerCP (BD Bioscience, San Jose, CA, USA), anti-CD25-APC (BD Bioscience), and anti-CD127-FITC (BD Bioscience) for 20 minutes in the dark at 4°C. Afterward, they were stained with anti-IL-17-PE (BD Bioscience) for 30 minutes in the dark at 4°C. Isotype antibodies were used to separate positive and negative cells in the PerCP, APC, FITC, and PE fluorescence channels. The samples were analyzed using an FACS Calibur analyzer (BD Bioscience), and data was acquired and analyzed using Cell Quest Pro Software (BD Bioscience) and Flow Jo software version 8. 6 (Tree Star Inc., Ashland, OR USA), respectively.

### ELISA for IL-35,IL-17, IL-4,TNF-α

To quantify cytokine production, we collected cell-free supernatants from cell cultures, and we used commercial ELISA kits (CusaBio, Wuhan, Hubei Province, China) to measure the concentration of cytokines according to the manufacturer’ s instructions.

### Statistical analysis

Continuous measurement data are reported as mean ± standard deviation (SD), while enumeration data are presented as n (%). To evaluate differences between groups, we used the rank-sum test and independent-samples t-test as appropriate. Additionally, we conducted Pearson and partial correlation analyses to evaluate correlations between parameters. A two-sided P-value less than 0. 05 was considered statistically significant. All data analyses were performed using SPSS version 22. 0 (SPSS Inc., Chicago, IL, USA).

## Results

### Serum biochemical indices

The study results indicated a significant increase in the serum levels of TBIL, ALT, and AST 48 hours after modeling in both the model group and the treatment groups compared to the control group (P < 0.05). Additionally, the levels of TBIL, ALT, and AST in the IL-35 plasmid group and dexamethasone group were significantly lower compared to the model group, while those in the anti-IL-35 antibody group were significantly higher than the model group (P < 0.05). However, no significant difference was observed between the IL-35 plasmid group and the dexamethasone group (P > 0.05) **(Table 1)**.

**Table 1.**
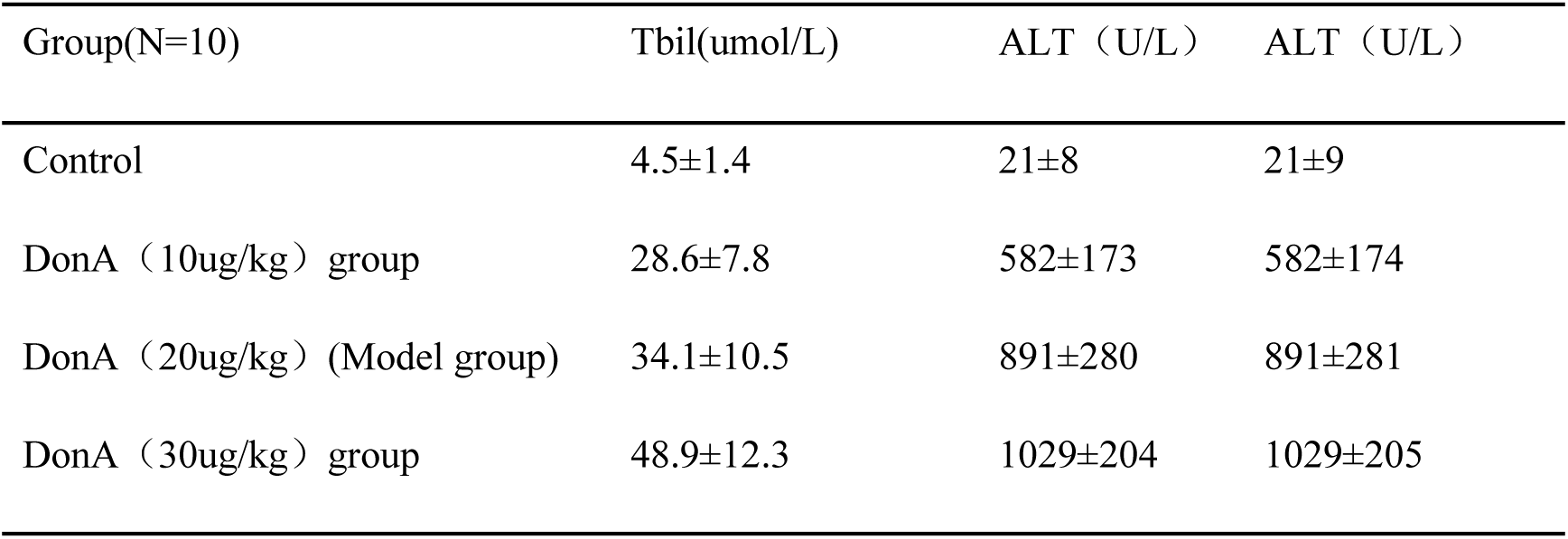
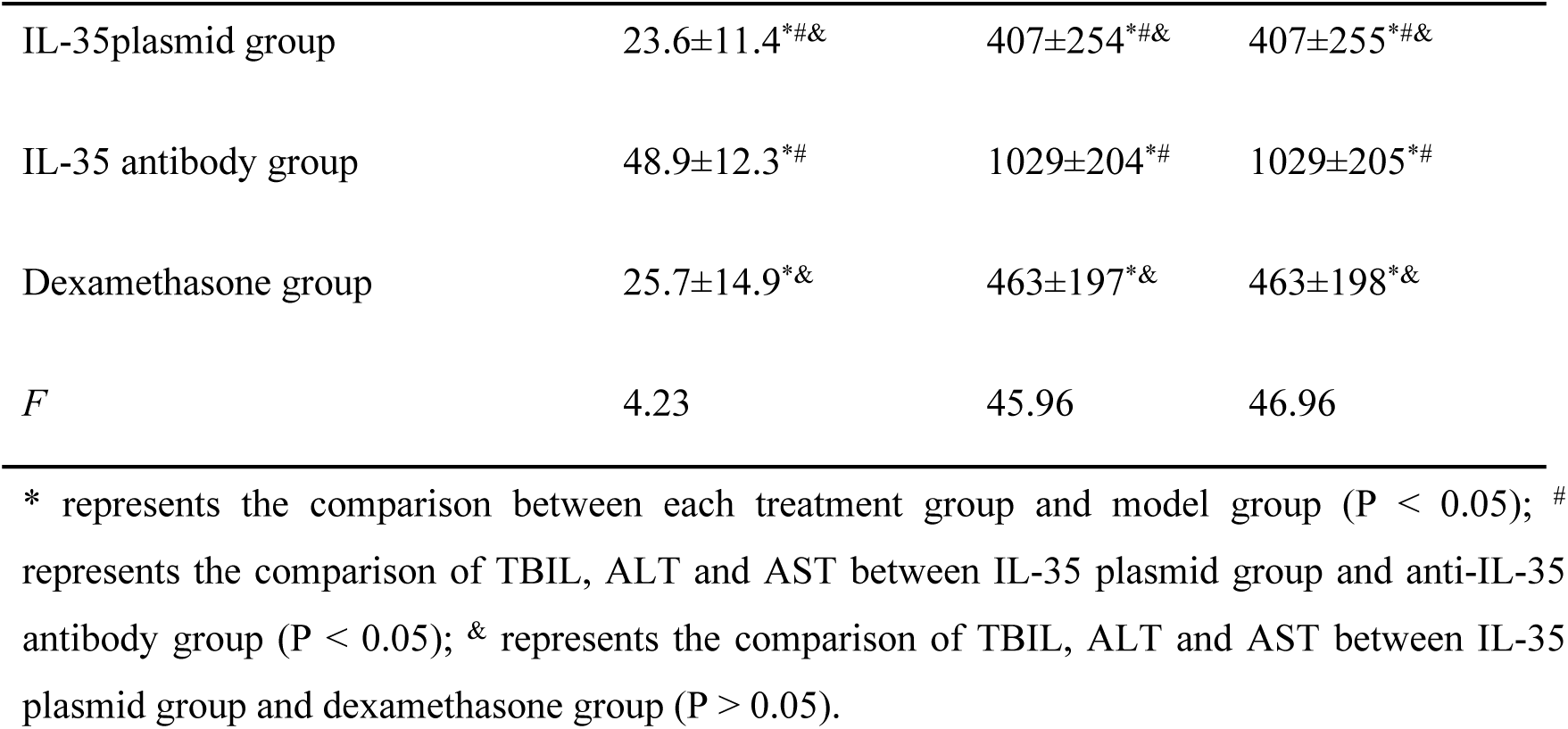
Levels and comparison of serum TBIL, ALT and AST of mice in each group.

| Group(N=10) | Tbil(umol/L) | ALT (U/L) | ALT (U/L) |
| --- | --- | --- | --- |
| Control | 4.5±1.4 | 21±8 | 21±9 |
| DonA (10ug/kg) group | 28.6±7.8 | 582±173 | 582±174 |
| DonA (20ug/kg) (Model group) | 34.1±10.5 | 891±280 | 891±281 |
| DonA (30ug/kg) group | 48.9±12.3 | 1029±204 | 1029±205 |
| IL-35plasmid group | 23.6±11.4*#& | 407±254*#& | 407±255*#& |
| IL-35 antibody group | 48.9±12.3*# | 1029±204*# | 1029±205*# |
| Dexamethasone group | 25.7±14.9*& | 463±197*& | 463±198*& |
| *F* | 4.23 | 45.96 | 46.96 |

\* represents the comparison between each treatment group and model group ( $P < 0.05$ ); # represents the comparison of TBIL, ALT and AST between IL-35 plasmid group and anti-IL-35 antibody group ( $P < 0.05$ ); & represents the comparison of TBIL, ALT and AST between IL-35 plasmid group and dexamethasone group ( $P > 0.05$ ).

### HE staining of mouse liver tissue

The control group exhibited the normal morphology of hepatocytes and hepatic lobule structure. In the low-dose group, we observed mild edema-like degeneration of some hepatocytes in the central vein of hepatic lobules with a small amount of monocytes and lymphocytes infiltrating around them, but no obvious necrosis was found. In the middle-dose group, we observed plate-like and debris-like necrotic lesions on the microscope. There was serious infiltration of lymphocytes and mononuclear macrophages in the portal area, consistent with the pathological changes in the early stage of liver failure. In the high-dose group, we observed diffuse degeneration and necrosis of hepatocytes with massive necrotic lesions on the microscope, consistent with the pathological diagnosis of liver failure. In the IL-35 plasmid group and dexamethasone group, we observed a small amount of lymphocyte infiltration around the central vein of the hepatic lobule with mild degeneration and edema of hepatocytes. However, we did not observe any obvious necrotic changes. The degree of inflammation and necrosis in these groups was significantly less than that in the model group and anti-IL-35 antibody group. In the anti-IL-35 antibody group, we observed more local hepatocyte degeneration and necrosis on the microscope, and the degree of necrosis was significantly higher than that in the other groups. Lymphocyte infiltration was also obvious (**Fig. 1)**.

**Fig. 1.**
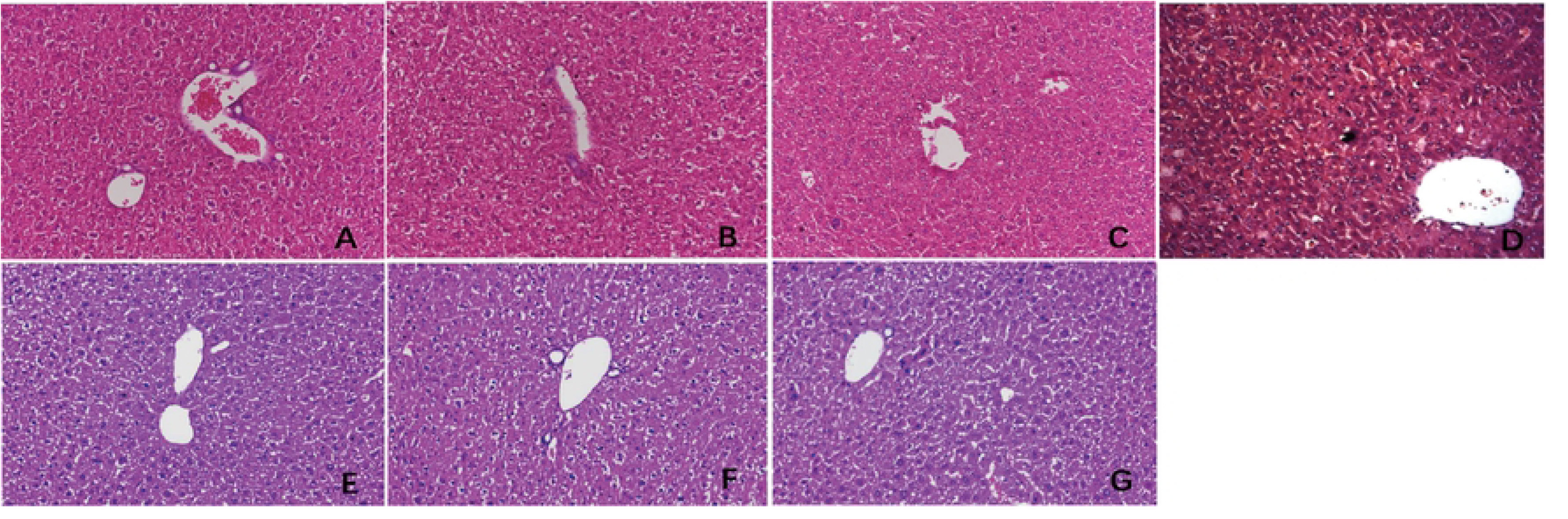
HE staining of liver tissue in control group, different doses of concanavalin and different intervention groups. (A) control group. (B) DonA (10ug/kg) group. (C) DonA (2Oug/kg) group. (D) DonA (30ug/kg) group. (E) IL-35pIasmid group. (F) Dexamethasone group. (G) anti-IL-35antibody group.

### Liver pathological index score (PI)

Compared to the control group, we observed a significant increase in the pathological index in the early stage of liver failure in the model group and every treatment group (P < 0.05). However, when compared to the model group, we found a significant decrease in the pathological index of the IL-35 plasmid group and dexamethasone group whereas there was a significant increase in the pathological index of the anti-IL-35 antibody group (P < 0. 05). Nevertheless, no significant difference was observed in the pathological index between the IL-35 plasmid group and the dexamethasone group (P > 0.05). These results suggest that IL-35 and dexamethasone may have protective effects on liver injury **(Table 2)**.

**Table 2.**
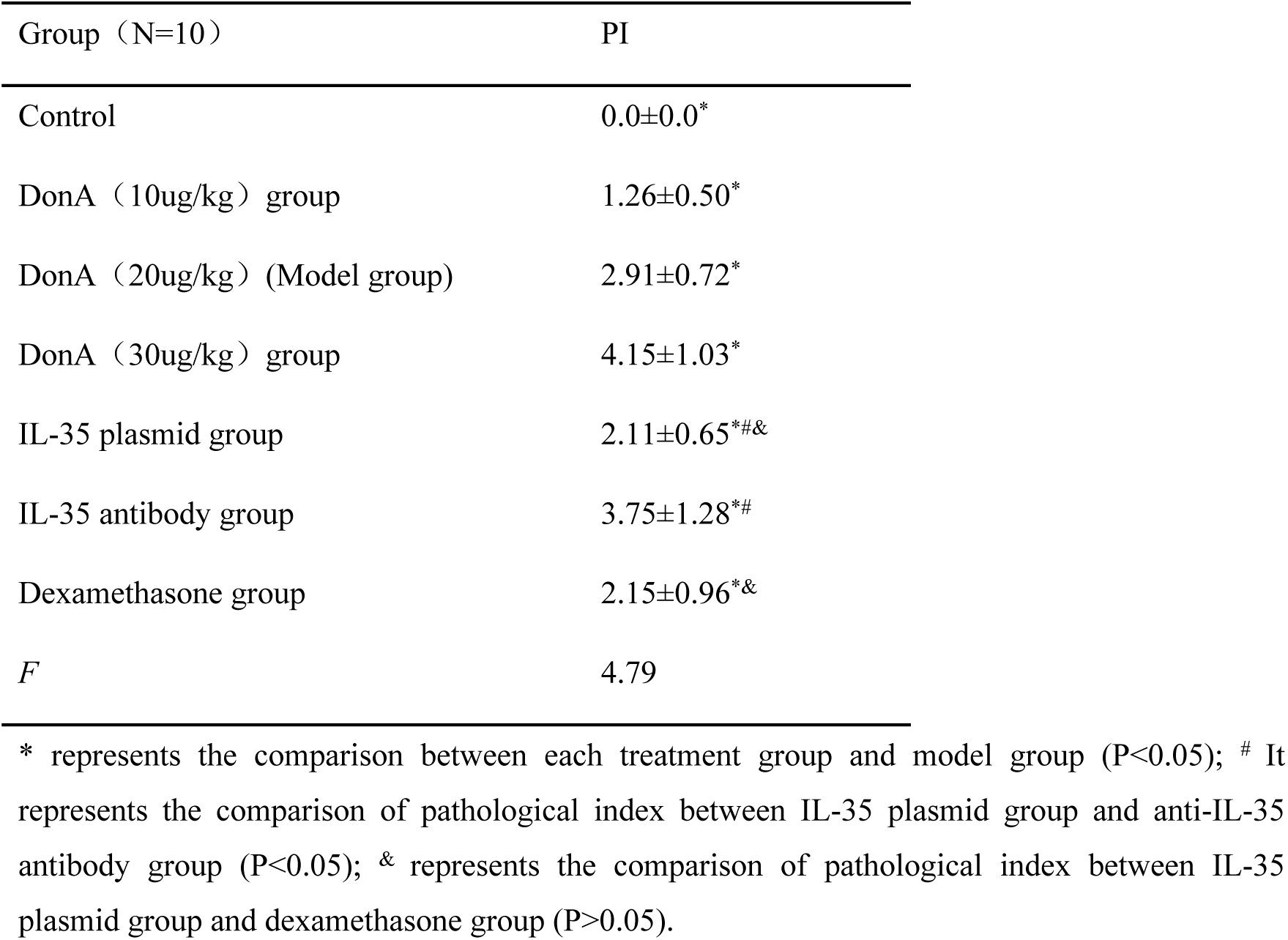
Levels and comparison of liver histopathological index (PI) of mice in each group.

### Serum IL-4, IL-17, IL-35 and TNF-α

We observed significantly higher levels of IL-35 in the IL-35 plasmid group and dexamethasone group when compared to the blank control group. When compared to the model group, we observed significantly lower levels of serum IL-4, IL-17, and TNF in the IL-35 plasmid group and dexamethasone group, while IL-35 levels were significantly higher than those in the model group and anti-IL-35 group. However, there was no significant difference between the IL-35 plasmid group and the dexamethasone group in terms of the levels of IL-4, IL-17, and TNF (P > 0. 05) **(Table 3, Fig. 2)**.

**Fig. 2.**
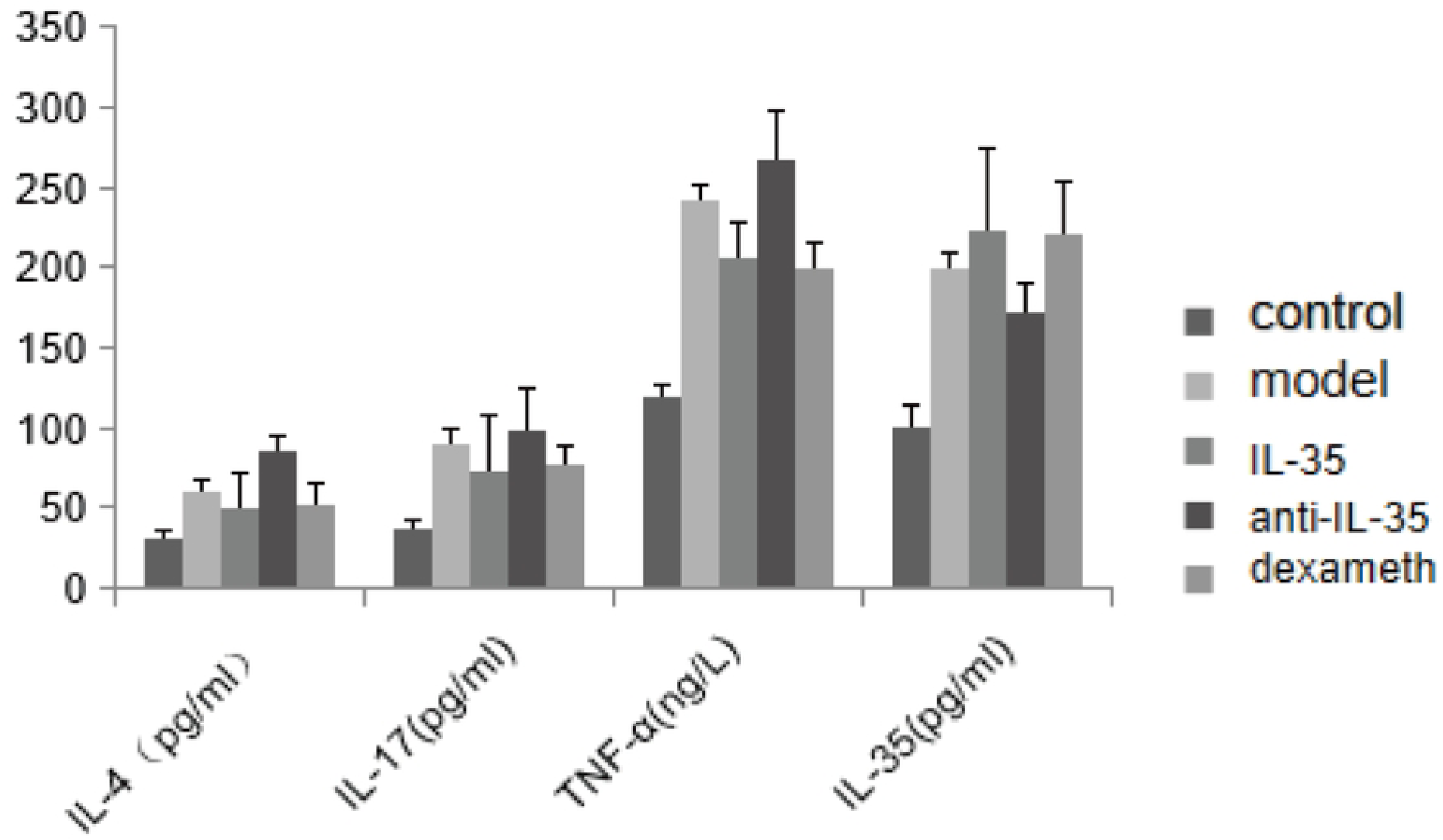
Serum IL-4, IL-17, IL-35 and TNF-α in blank control group, model group and each treatment group.

**Table 3.**
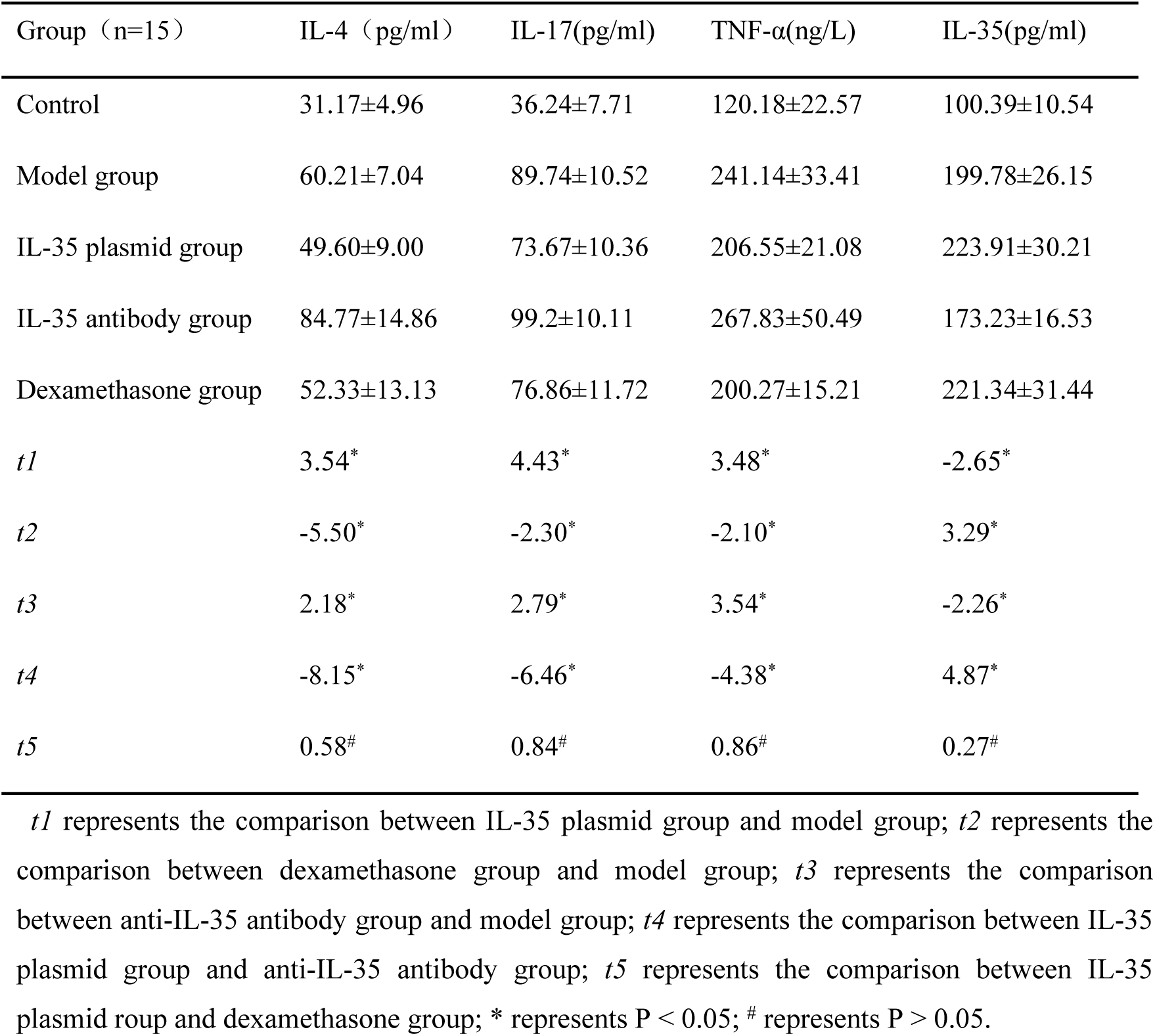
Serum IL-4, IL-17, IL-35 and TNF-α in control group, model group and each treatment group.

| Group (n=15) | IL-4 (pg/ml) | IL-17(pg/ml) | TNF-α(ng/L) | IL-35(pg/ml) |
| --- | --- | --- | --- | --- |
| Control | 31.17±4.96 | 36.24±7.71 | 120.18±22.57 | 100.39±10.54 |
| Model group | 60.21±7.04 | 89.74±10.52 | 241.14±33.41 | 199.78±26.15 |
| IL-35 plasmid group | 49.60±9.00 | 73.67±10.36 | 206.55±21.08 | 223.91±30.21 |
| IL-35 antibody group | 84.77±14.86 | 99.2±10.11 | 267.83±50.49 | 173.23±16.53 |
| Dexamethasone group | 52.33±13.13 | 76.86±11.72 | 200.27±15.21 | 221.34±31.44 |
| *t1* | 3.54* | 4.43* | 3.48* | -2.65* |
| *t2* | -5.50* | -2.30* | -2.10* | 3.29* |
| *t3* | 2.18* | 2.79* | 3.54* | -2.26* |
| *t4* | -8.15* | -6.46* | -4.38* | 4.87* |
| *t5* | 0.58<sup>#</sup> | 0.84<sup>#</sup> | 0.86<sup>#</sup> | 0.27<sup>#</sup> |

*t1* represents the comparison between IL-35 plasmid group and model group; *t2* represents the comparison between dexamethasone group and model group; *t3* represents the comparison between anti-IL-35 antibody group and model group; *t4* represents the comparison between IL-35 plasmid group and anti-IL-35 antibody group; *t5* represents the comparison between IL-35 plasmid group and dexamethasone group; \* represents $P < 0.05$ ; # represents $P > 0.05$ .

### Comparison of immunohistochemical and immunofluorescence results of IL-35 and IL-17 in liver tissue of mice

The results of the immunohistochemistry analysis revealed clear structures of positive cells, with IL-12a (IL-35 subunit) and IL-17 appearing as brown-yellow particles showing good positioning, and the degree of staining was significantly higher than the background color. Further analysis through fluorescence staining showed that Foxp3 was the characteristic staining of regulatory T cells with a green fluorescence staining color, and IL-12a (IL-35) appeared as red fluorescence. Our results confirmed that IL-35 is mainly secreted by regulatory T cells and that there was higher expression of IL-35 observed in Foxp3 positive regulatory T cells (**Fig. 3**, **Fig. 4)**.

**Fig. 3.**
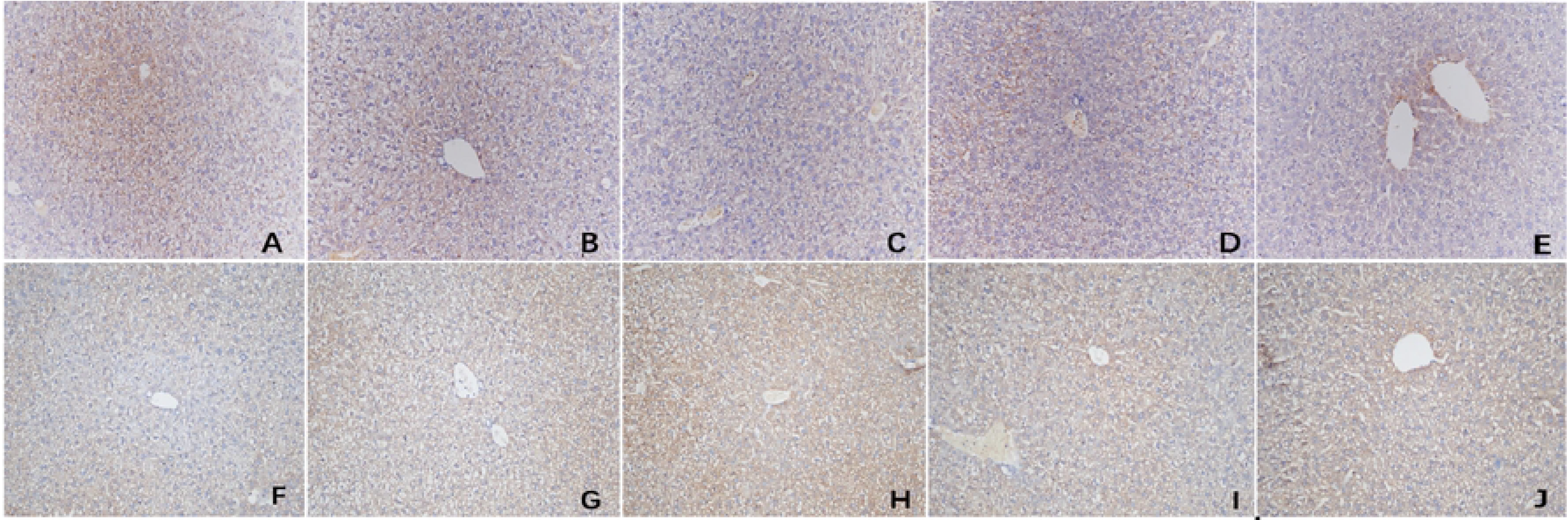
Comparison of immunohistochemical results of IL-35 and IL-17 in liver tissue of mice in control group, PLF model group and each treatment group. (A) IL-17 in liver of control group mice. (B) IL-17 in liver of model group mice. (C) IL-17 in liver of IL-35 plasmid group mice. (D) IL-17 in liver of anti-IL-35 group mice. (E) IL-17 in liver of dexamethasone group mice. (F) IL-35 in liver of control group mice. (G) IL-35 in liver of model group mice. (H) IL-35 in liver of IL-35 plasmid group mice. (I) IL-35 in liver of anti -IL-35 group mice. (J) IL-35 in liver of dexamethasone group mice.

**Fig. 4.**
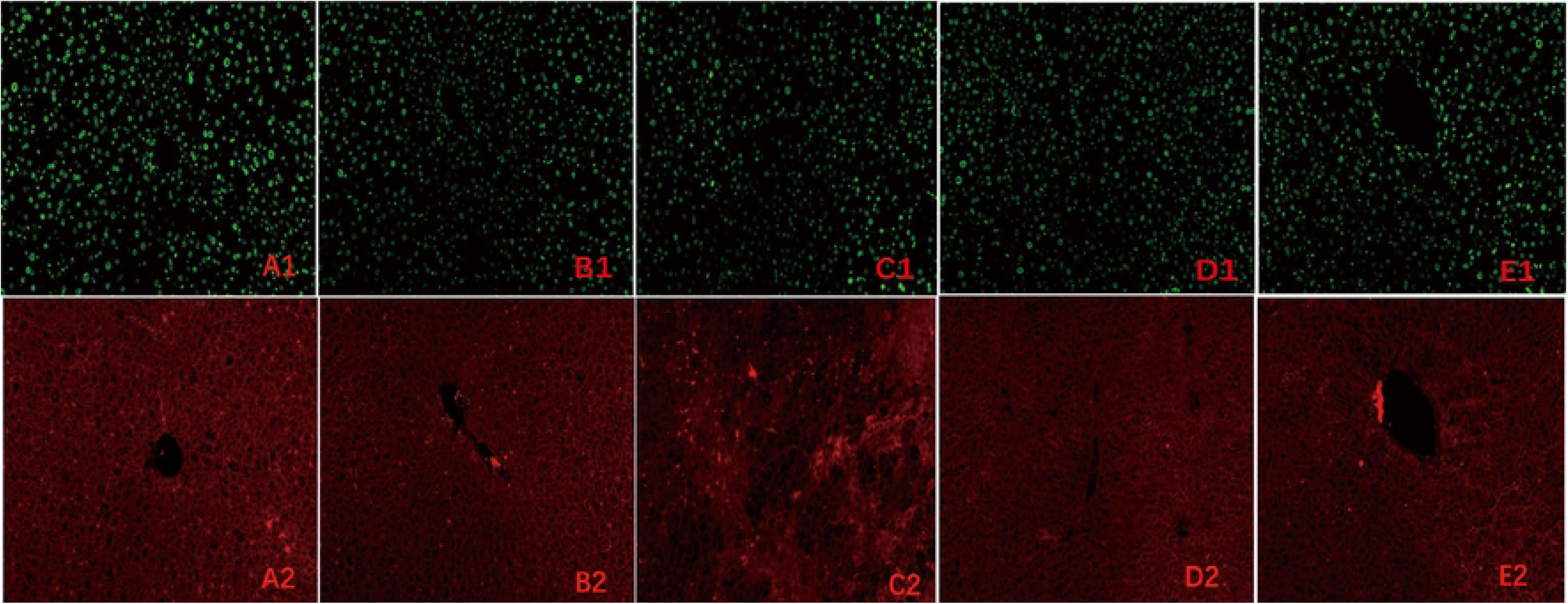
Comparison of immunofluorescence results of IL-35 and FOXP3 in liver tissue of mice in control group, PLF model group and each treatment group. (A1)FOXP3 in liver of control group mice. (A2) IL-35 in liver of control group mice. (Bl) FOXP3 in liver of model group mice. (B2)IL-35 in liver of model group mice. (Cl) FOXP3 in liver of IL-35 plasmid group mice. (C2) IL-35 in liver of IL-35 plasmid group mice. (DI) FOXP3 in liver of anti-IL-35 group mice. (D2) IL-35 in liver of anti-IL-35 group mice. (El) FOXP3 in liver of dexamethasone group mice. (E2) IL-35 in liver of dexamethasone group mice.

### The positive areas of IL-17 and IL-35 in the liver tissue of mice

We observed significantly higher positive areas of IL-17 and IL-35 in the liver tissue of mice in the PLF model group and each treatment group compared to the control group. However, we found that the positive areas of IL-17 in the IL-35 plasmid group and dexamethasone group were significantly lower, while IL-35 was higher than that in the PLF model group. Conversely, in the anti-IL-35 antibody group, we observed a significantly higher positive area of IL-17 while IL-35 was lower than that in the PLF model group. Additionally, we observed a significantly lower positive area of IL-17 in the IL-35 plasmid group, while IL-35 was higher than that in the dexamethasone group (P < 0.05). However, we did not observe any significant differences between the IL-35 plasmid group and dexamethasone group in the peripheral serum (ELISA) (P > 0. 05) (**Table 4**, **Fig. 5-6)**.

**Fig. 5.**
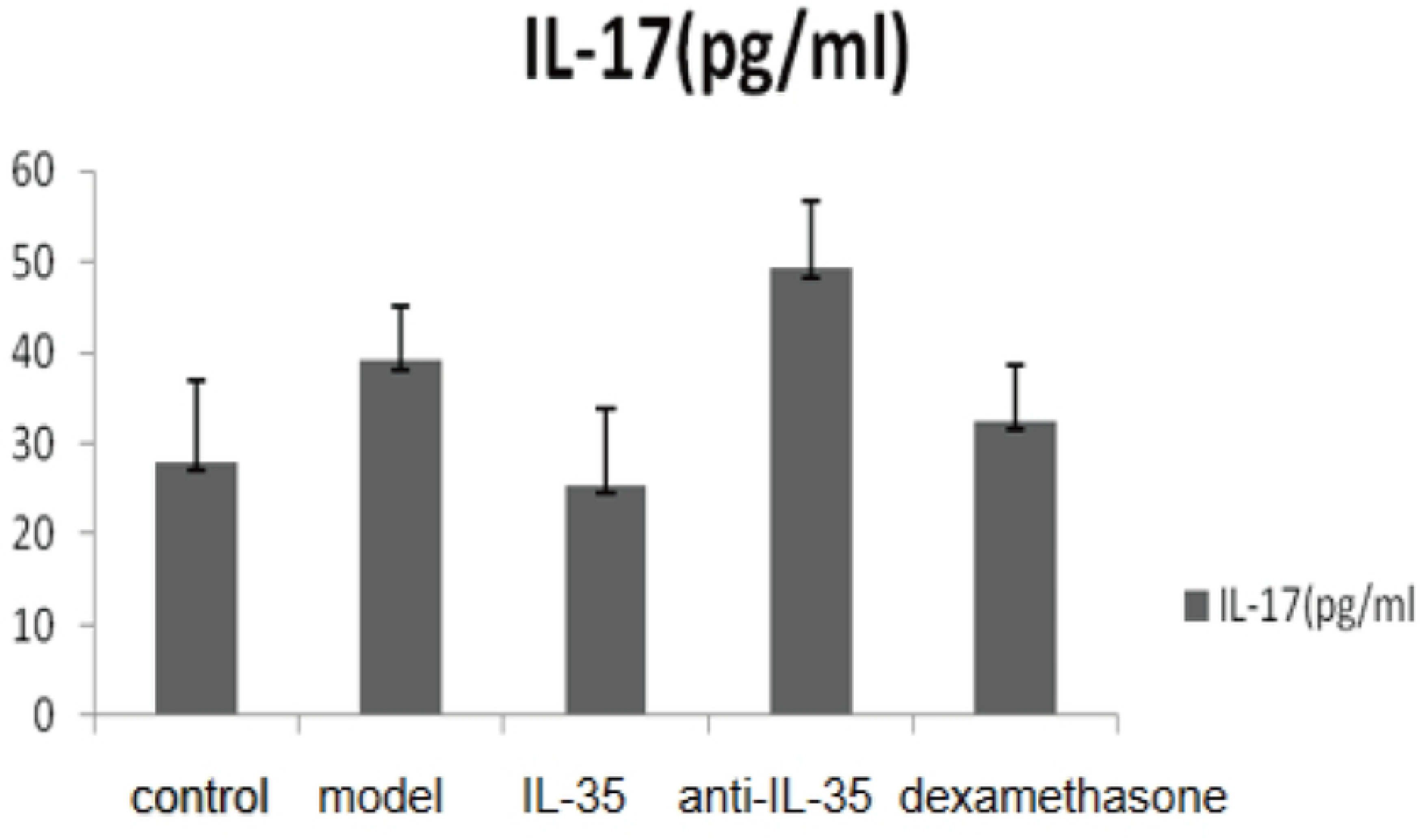
Positive area of IL-17 immunohistochemistry in liver tissue of mice.

**Fig. 6.**
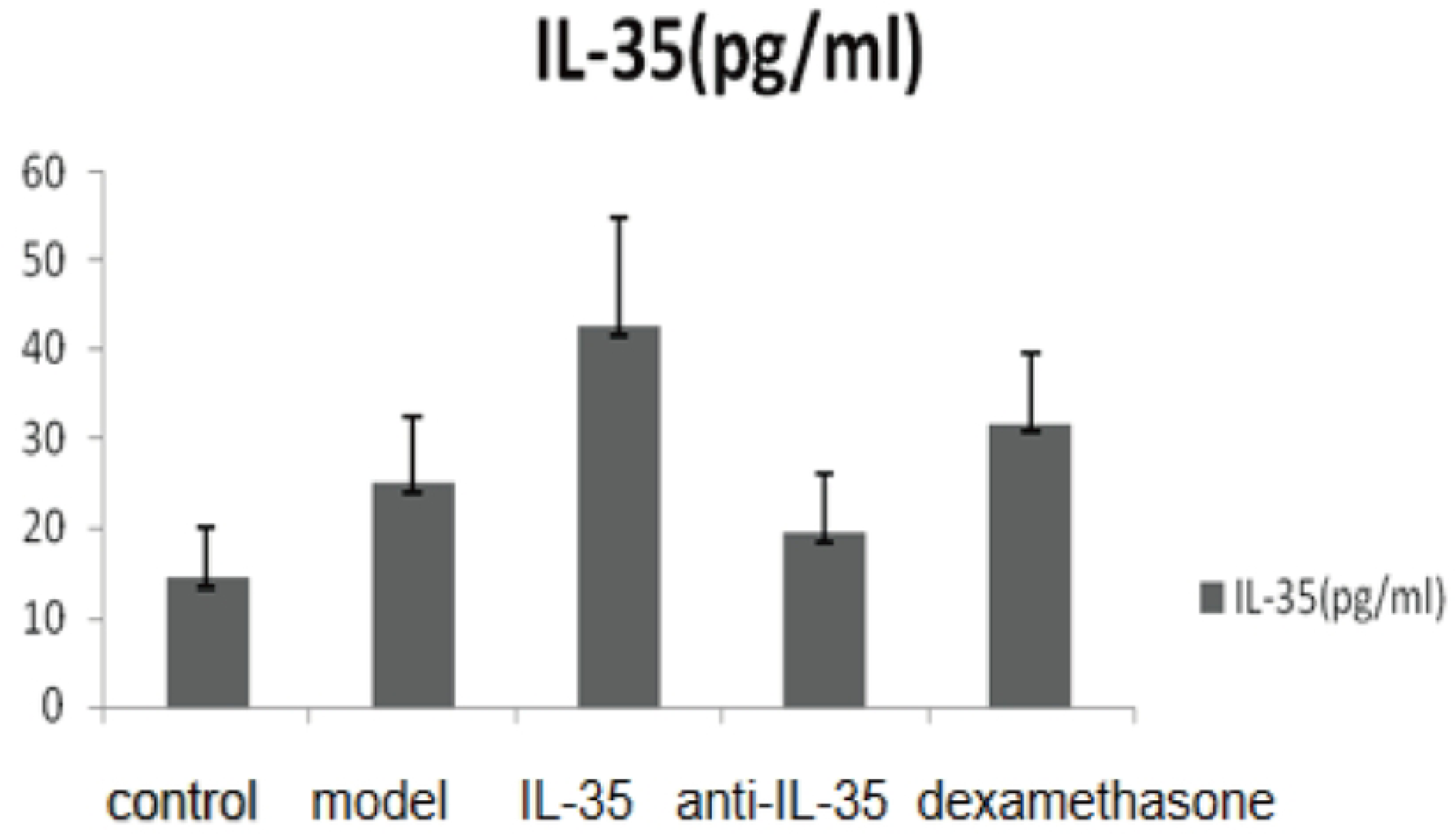
Positive area of IL-35 immunohistochemistry in liver tissue of mice.

**Table 4.** Comparison of positive expression areas of IL-17 and IL-35 in liver tissue of mice in control group, model group and each treatment group.

| Group (n=15) | IL-17(pg/ml) | IL-35(pg/ml) |
| --- | --- | --- |
| Control | 28.07±8.96 | 14.26±5.86 |
| Model group | 39.12±6.13 | 24.95±7.48 |
| IL-35 plasmid group | 25.43±8.28 | 42.63±12.18 |
| IL-35 antibody group | 49.31±7.5 | 19.39±6.59 |
| Dexamethasone group | 32.43±6.28 | 31.67±7.81 |
| *t1* | 5.22* | 4.79* |
| *t2* | 3.46* | 2.49* |
| *t3* | 2.58* | 2.28* |
| *t4* | 2.45* | 2.54* |

*t1* represents the comparison between IL-35 plasmid group and model group; *t2* represents the comparison between anti-IL-35 antibody and model group, dexamethasone group and model group; *t3* represents the comparison between dexamethasone group and model group; *t4* represents the comparison between IL-35 plasmid group and dexamethasone group; \* represents $P < 0.05$ .

### Laboratory examination of patients with HBV-PLF at baseline after admission and one week after glucocorticoid treatment

After one week of methylprednisolone (1mg/kg) pulse therapy, we observed that 18 out of 22 patients showed improvement while 4 patients still progressed to liver failure. Among these, 3 patients showed improvement after receiving active symptomatic support and plasma exchange treatment, while one patient was transferred to another hospital for liver transplantation due to liver cirrhosis and hepatic encephalopathy. Our analysis showed that there was no significant difference in age and gender between the patient group and the healthy control group. Additionally, we found that the activities of total bilirubin, alanine aminotransferase, aspartate aminotransferase, and prothrombin were significantly lower than those before glucocorticoid treatment, both at admission and one week after glucocorticoid treatment. Based on these findings, we can conclude that the liver injury of the patients was relieved after glucocorticoid treatment(**Table 5)**.

**Table 5.**
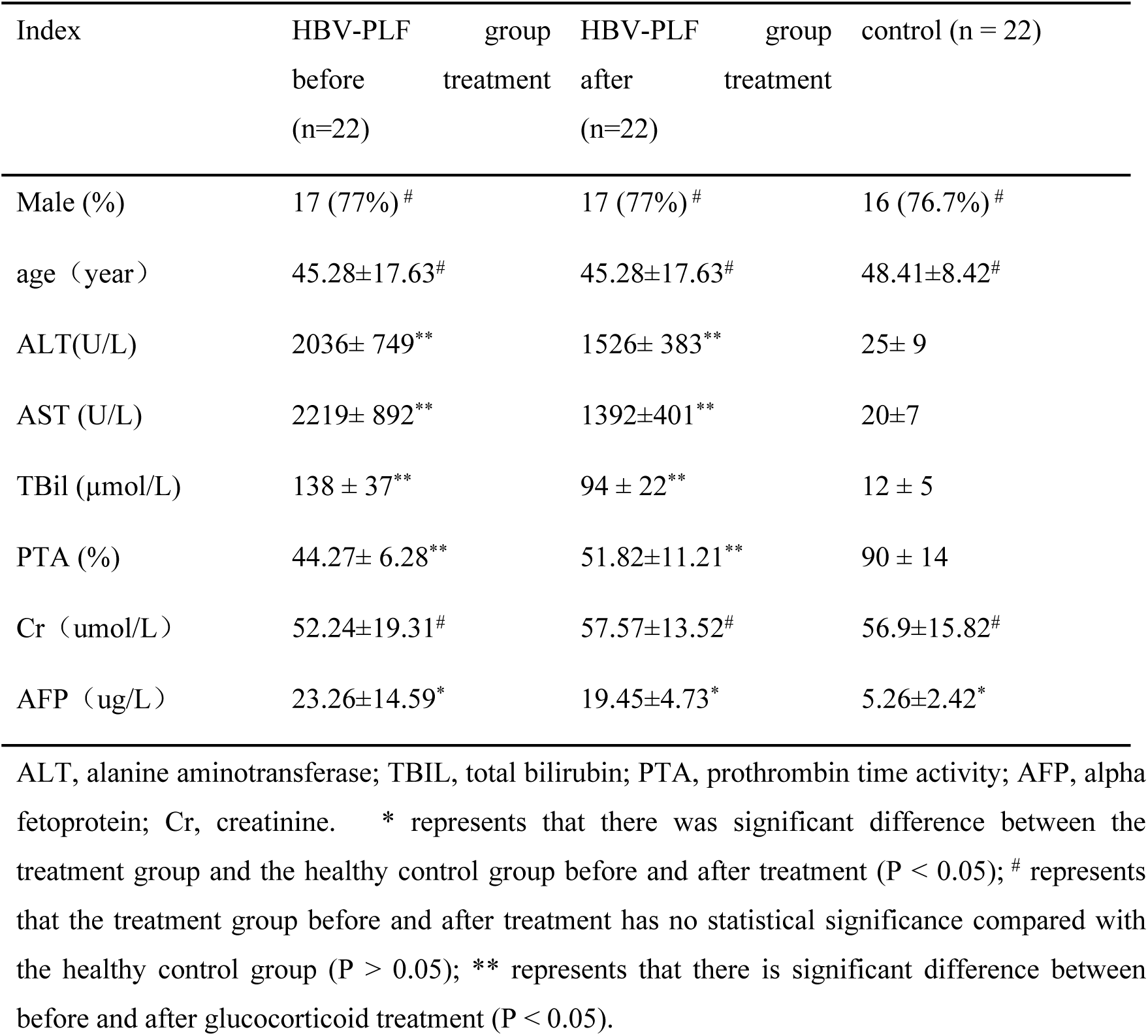
Comparison of liver function indexes, Cr and AFP before and after glucocorticoid therapy in patients with HBV-PLF.

### Treg and Th17 of HBV-PLF patients were detected by Flow cytometry at baseline and one week after glucocorticoid treatment

Our analysis showed that before glucocorticoid treatment, Tregs, especially Th17 in HBV-PLF patients, were significantly higher than those in the healthy group. This resulted in a significantly lower Treg/Th17 ratio. However, after one week of glucocorticoid treatment, we found that Tregs were significantly higher while Th17 were significantly lower, and Treg/ Th17 was significantly higher than that before treatment. These findings suggest that glucocorticoid treatment may contribute to restoring an appropriate balance between Tregs and Th17 in HBV-PLF patients (**Table 6)**.

**Table 6.** Treg and Th17 of HBV-PLF patients were detected by Flow cytometry at baseline and one week after glucocorticoid treatment.

| Group | Treg (%) | Th17 (%) | Treg/Th17 |
| --- | --- | --- | --- |
| HBV-PLF (before) | 5.38±0.82 | 2.29±0.50 | 2.50±0.88 |
| HBV-PLF (after) | 5.86±0.90 | 1.80±0.50 | 3.65±1.80 |
| control | 4.03±0.71 | 0.33±1.0 | 13.18±4.61 |
| $t_1$ | 6.09* | 7.37* | 4.59* |
| $t_2$ | 6.28* | 18.95* | 10.54* |

$t_1$ represents paired t-test before and after treatment; $t_2$ represents paired t-test compared with healthy control group before treatment; \* represents $P < 0.01$ .

### Serum IL-35 and IL-17 in HBV-PLF patients by ELISA and healthy control group before and one week after glucocorticoid treatment

Our analysis showed that the levels of IL-35 and IL-17 were significantly higher in HBV-PLF patients than those in the healthy group before glucocorticoid treatment. However, after one week of glucocorticoid treatment, we observed that the level of IL-35 was significantly higher, while IL-17 was significantly lower than that before the intervention. These changes were consistent with the alterations observed in Treg and Th17 as determined by flow cytometry. These results suggest that the immunomodulatory effects of glucocorticoid treatment may contribute to the regulation of IL-35 and IL-17 levels in HBV-PLF patients (**Table 7)**.

**Table 7.** Serum IL-35 and IL-17 in HBV-PLF patiens by ELISA and healthy control group before and one week after glucocorticoid treatment.

| Group | IL-35 (pg/ml) | IL-17 (pg/ml) |
| --- | --- | --- |
| HBV-PLF (before) | 632.59±73.26 | 68.18±11.39 |
| HBV-PLF (after) | 701.95±81.83 | 59.05±8.74 |
| control | 61.32±7.79 | 33.14±4.78 |
| $t_1$ | 6.39* | 7.63* |
| $t_2$ | 34.72* | 12.83* |

$t_1$ represents paired t-test before and after treatment; $t_2$ represents paired t-test compared with healthy control group before treatment; \* represents $P < 0.01$ .

### Correlation between serum IL-35 level and Treg / Th17 ratio in HBV-PLF patients

The findings suggest that there was a significant positive correlation between IL-35 and Treg/ Th17 (r = 0.69). However, no significant correlation was observed between IL-17 and Treg/Th17(**Fig.7-8)**.

**Fig. 7.**
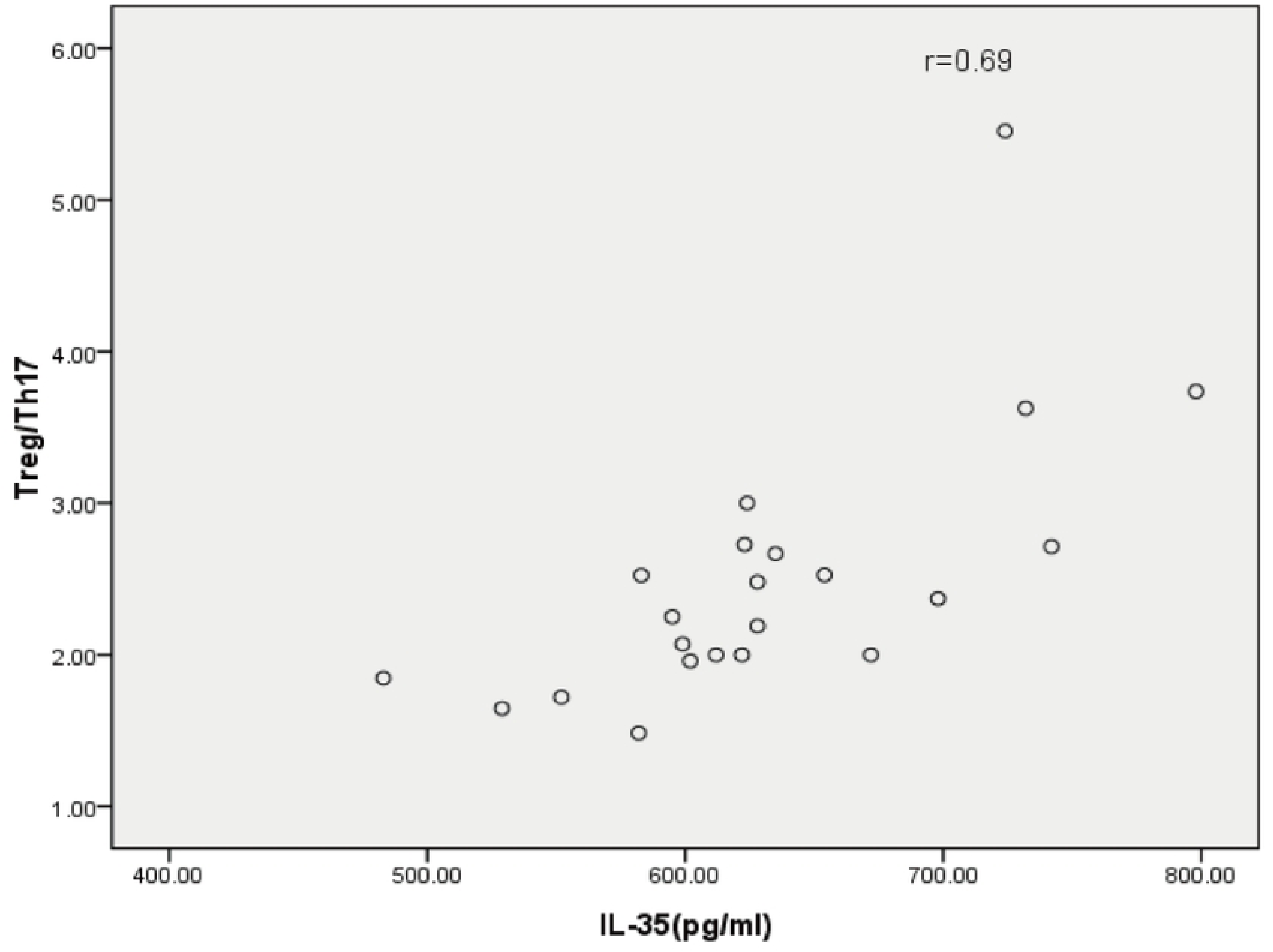
Correlation between serum IL-35and Treg / Th 17 ratio in HBV-PLF Patients with pre hepatitis B related liver failure.

**Fig. 8.**
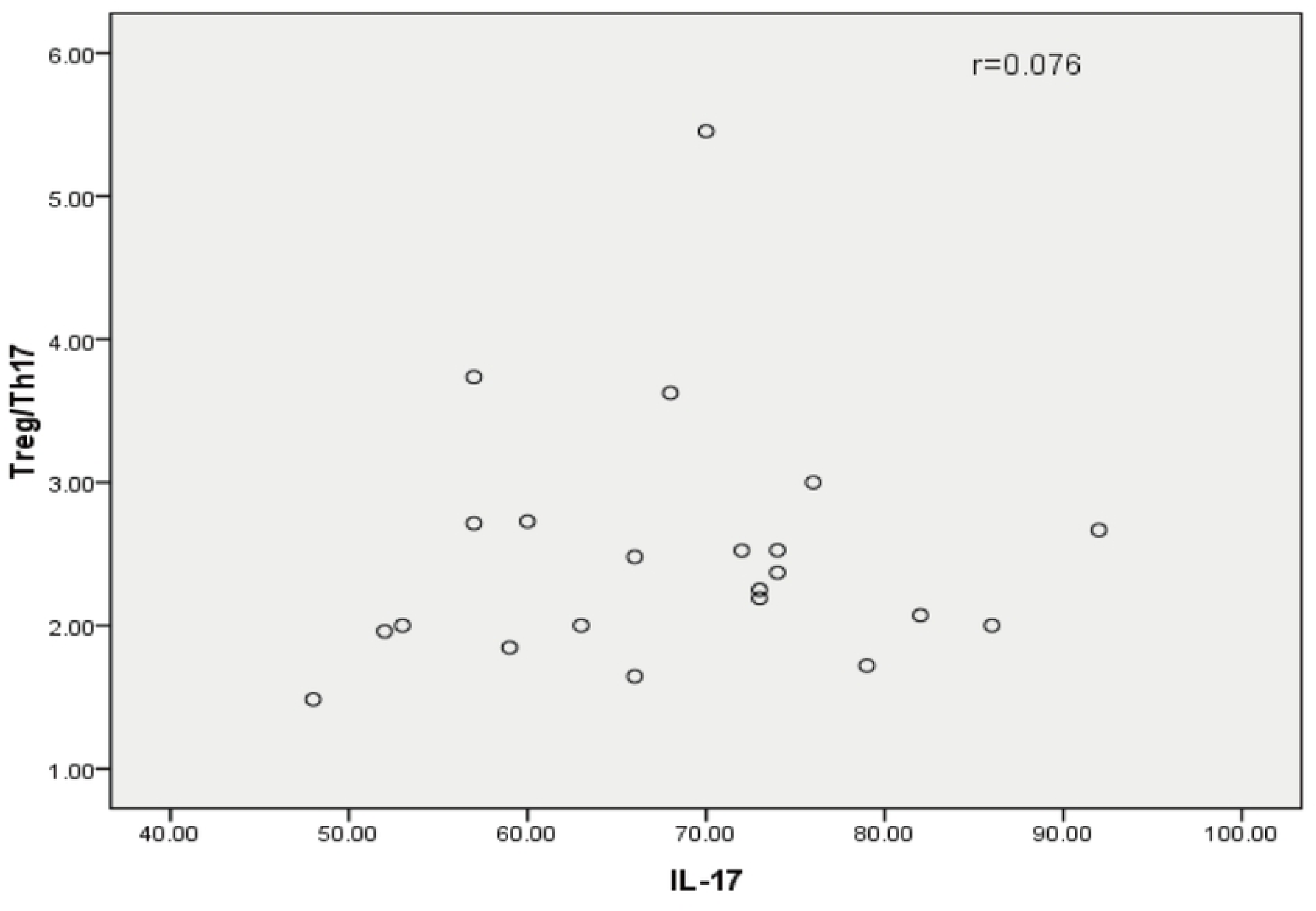
Correlation between serum IL-17 and Treg / Thl7 ratio in HBV-PLF patients with pre hepatitis B related liver failure.

## Discussion

In summary, Hepatitis B virus (HBV) is a common cause of severe hepatitis in China, and immune disorder and liver cell necrosis are key mechanisms responsible for hepatitis B-related liver failure. Cellular immunity, including T cells, Th17 cells, NK/NK T cells, monocyte macrophages, etc., is primarily responsible for causing hepatocyte necrosis. The proportion and functional changes of Treg/Th17 and the levels of related cytokines, including IL-35 and IL-17, have been confirmed that can influence the body’s immune response in numerous studies[10–14]. IL-35 is mainly secreted by regulatory T cells (Treg cells) and plays a crucial role in regulating immune balance by altering the ratio of Th17/Treg and suppressing the positive immunity of Th17[15–18]. Glucocorticoids (GCS) can act as immunomodulatory agents by regulating the balance of Treg/Th17 and related cytokines[19]. Additionally, studies have shown that glucocorticoids can stimulate the production of IL-35 and boost the number of Treg cells by activating related signaling pathways[20].

The treatment of liver failure is a significant challenge in clinical practice. Early intervention and control of the condition may be key to successful treatment. Therefore, this study focuses on animal models and patients in the early stage of liver failure. Mice were divided into several groups, including blank, model, IL-35 plasmid, anti-IL-35 antibody, and dexamethasone groups. Our preliminary results suggest that the total bilirubin, ALT, and AST levels of mice in the plasmid and hormone groups were significantly lower than those in the anti-IL-35 antibody group, indicating that IL-35 and glucocorticoids provide a certain protective effect on hepatocytes in the prophase of liver failure. Furthermore, the ELISA results revealed that following the intervention of IL-35 plasmid, hormone, and anti-IL-35 antibody, the levels of IL-4, IL-17, and TNF-α in the plasmid and hormone groups were significantly lower than those observed in the model group. This finding further confirms the inhibitory effects of IL-35 and glucocorticoids on positive inflammatory factors in the model group.

To further validate our findings, we examined the pathological changes in the liver, which is considered the gold standard for assessing liver injury. Our results indicate that the liver damage area and pathological index in the IL-35 plasmid and dexamethasone groups are significantly lower than in the model and anti-IL-35 groups. This finding further confirms that IL-35 is involved in liver protection, which is consistent with research conclusions on IL-35 in HBV-related liver diseases[21]. The observed inhibitory effects of IL-35 and glucocorticoids on positive inflammatory factors and their protective effects on hepatocytes in the prophase of liver failure suggest that IL-35 and glucocorticoid-based treatments may be potential therapeutic approaches for treating early liver failure. However, further studies are needed to determine the safety and effectiveness of these treatments.

In addition to assessing the pathological changes, we also used immunohistochemistry and fluorescence staining to validate our findings. Our results indicate that under the intervention of IL-35 and dexamethasone, the expression of negative immune regulatory cells and cytokines, such as Treg cells and IL-35, was up-regulated, while positive immune cells and cytokines, such as Th17 cells and IL-17, were effectively controlled. Interestingly, when comparing the immunohistochemical positive areas of IL-17 and the negative immune inflammatory factor IL-35 in each group, we found that the positive area of IL-35 in the IL-35 plasmid group was significantly higher and IL-17 was lower than in the dexamethasone group. This suggests that IL-35 may have more advantages than glucocorticoids in inhibiting the expression of IL-17 in liver tissue. However, there was no significant difference in the level of IL-35 between the plasmid and dexamethasone groups in peripheral serum. This may indicate the existence of differences between peripheral immunity and local immunity in the liver tissues of mice in the prophase of liver failure. In conclusion, our findings confirm the inhibitory effects of IL-35 and glucocorticoids on positive immune factors and the enhancement of negative immune factors in liver injury, especially in the prophase of liver failure. Th17 cells play a crucial role in HBV-related hepatocyte immune injury by primarily secreting cytokines such as IL-17, IL-22, and IL-23. There is a positive correlation between Th17 cells and hepatocyte immune injury caused by HBV infection. At the same time, Treg cells inhibit the immune inflammatory reaction that has occurred. The Treg/ Th17 ratio is relatively balanced in thriving immune states. However, once this balance is disrupted, it may cause severe liver immune injury and even liver failure [22–25].

We also conducted a clinical study to evaluate the effect of glucocorticoids on immune regulation in patients in the prophase of liver failure. Our findings indicate that one week after glucocorticoid treatment, the levels of IL-35 and Treg cells in patients significantly increased compared with before treatment. In contrast, Th17 cells and IL-17 decreased significantly compared with before treatment. More importantly, the Treg/ Th17 ratio increased significantly compared with before treatment, although it remained slightly lower than that in healthy individuals. These results confirm the clinical significance of glucocorticoids in immune regulation in patients in the prophase of liver failure and in maintaining the stability of the Treg/Th17 ratio. Our results suggest that glucocorticoids may help to maintain the cellular and humoral immune balance and prevent the occurrence of liver failure in patients with early-stage hepatitis B-related liver failure by increasing the Treg/Th17 ratio and levels of IL-35.

It is important to note that our clinical study has some limitations. Firstly, due to the clinical treatment needs of patients, we could not establish a control group that received non-glucocorticoid treatment, and the study design may not have been rigorous enough. Secondly, the study time was relatively short, and the number of patients in the prophase of HBV-related liver failure was relatively small. Additionally, the number of cases with poor treatment outcomes after glucocorticoid treatment was also very few, which made it unsuitable to compare statistically between the hormone effective group and the ineffective group. Therefore, more long-term, randomized, controlled clinical trials with a larger sample size are needed to further explore the clinical value of glucocorticoids in immune regulation in patients with early-stage liver failure.

Absolutely, expanding the sample size and conducting more long-term, randomized, controlled clinical trials with a more rigorous study design will be critical in future research to establish more robust conclusions and better guide the immunomodulatory treatment of patients with early-stage liver failure. These efforts will ultimately help improve clinical outcomes for patients suffering from liver failure, which is a significant public health concern globally.

## Notes

### Competing Interest Statement

The authors have declared no competing interest.

